# Transposable elements generate regulatory novelty in a tissue-specific fashion

**DOI:** 10.1101/268771

**Authors:** Marco Trizzino, Aurélie Kapusta, Christopher D. Brown

## Abstract

**Background:** Transposable elements (TE) are an important source of evolutionary novelty in gene regulation. However, the mechanisms by which TEs contribute to gene expression are largely uncharacterized.

**Results:** Here, we leverage Roadmap and GTEx data to investigate the association of TEs with active and repressed chromatin in 24 tissues. We find 112 human TE types enriched in active regions of the genome across tissues. SINEs and DNA transposons are the most frequently enriched classes, while LTRs are often enriched in a tissue-specific manner. We report across-tissue variability in TE enrichment in active regions. Genes with consistent expression across tissues are less likely to be associated with TE insertions. TE presence in repressed regions similarly follows tissue-specific patterns. Moreover, different TE classes correlate with different repressive marks: Long Terminal Repeat Retrotransposons (LTRs) and Long Interspersed Nuclear Elements (LINEs) are overrepresented in regions marked by H3K9me3, while the other TEs are more likely to overlap regions with H3K27me3. Young TEs are typically enriched in repressed regions and depleted in active regions. We detect multiple instances of TEs that are enriched in tissue-specific active regulatory regions. Such TEs contain binding sites for transcription factors that are master regulators for the given tissue. These TEs are enriched in intronic enhancers, and their tissue-specific enrichment correlates with tissue-specific variations in the expression of the nearest genes.

**Conclusions:** We provide an integrated overview of the contribution of TEs to human gene regulation. Expanding previous analyses, we demonstrate that TEs can potentially contribute to the turnover of regulatory sequences in a tissue-specific fashion.

## Background

Sequences derived from transposable element (TE) insertions make up roughly half of the length of the human genome. Several TE groups still show transposing activity in humans, including Long Terminal Repeat Retrotransposons (mostly ERV1-LTRs; [1–3]), Long Interspersed Nuclear Elements (LINEs, mostly L1s; [4–5]), Short Interspersed Nuclear Elements (SINEs) of the Alu families [6,7], and SINE-VNTR-*Alus* (SVAs; [8,9]).

Multiple elegant studies have demonstrated that TE sequences play a functional role in eukaryotic gene regulation [10–32]. Consistently, we recently demonstrated that TEs are the primary source of evolutionary novelty in primate gene regulation, and reported that the large majority of newly evolved human and ape specific liver cis-regulatory elements are derived from TE insertions [33]. Similarly, other studies have shown that the recruitment of novel regulatory networks in the uterus was likely mediated by ancient mammalian TEs [21,22], and that TEs have a role in pluripotency [34]. Conversely, other researchers have proposed that TE exaptation into regulatory regions is rare [35], and that TE silencing may not be a major driver of regulatory evolution in primates [36].

Given these contrasting lines of evidence, we aimed to shed light on the contribution of TEs to the evolution of the tissue-specific regulation of human gene expression. For this purpose, we took advantage of publicly available data [37,38] to investigate patterns of TE overlap with tissue-specific histone modification states and to characterize the contribution of TEs to tissue-specific gene expression. We find that a significant fraction of the existing human TEs are enriched in regions of the genome bearing epigenetic hallmarks of active or repressed chromatin, suggesting they could potentially be actively regulated by the cellular machinery. DNA transposons and SINEs represent the most frequently enriched classes across tissues, while LTR-ERV1s are the TEs that more commonly show tissue-specific enrichment and active regulatory activity. TE enrichment in active and repressed chromatin exhibits tissue-specific patterns. Genes with consistent expression across tissues are less likely to be associated with a local TE insertion. We detect multiple instances of TEs showing tissue-specific enrichment in active and repressed regions, and demonstrate that they contain binding sites for transcription factors that are tissue-specific master regulators.

## Results

### Specific TE families are enriched in active and repressed genomic regions

To investigate the extent to which TEs contribute to the regulation of human gene expression, we leveraged publicly available data from the Roadmap Epigenomics Project [37] and from the GTEx Project [38]. We focused on 24 primary tissues and cell types that were processed by both consortia (Supplementary Table S1). Using five different histone modifications (H3K4me1, H3K4me3, H3K36me3, H3K9me3, and H3K27me3), Roadmap segmented the human genome into 15 regulatory classes, reflecting different degrees and types of regulatory activity. We took advantage of this classification to define active (H3K4me1, H3K4me3, H3K36me3) and repressed (H3K9me3, and H3K27me3) chromatin regions in each of the studied tissues.

To test for TE enrichment in active and repressed chromatin, we used the TE-Analysis pipeline ([39]; https://github.com/4ureliek/TEanalysis; Supplemental File S1). This pipeline is designed to output the TE composition of given features, such as TE counts and TE amounts, aiming to detect potential TE enrichments in the select features. As expected, we find that the majority of human TEs are significantly depleted from regions marked as active by Roadmap histone modifications (mean 83.9% of TEs; FDR <5%; Supplementary Table S2). Nevertheless, 112 TE families (9.07% of the annotated TE families in the human genome) are significantly enriched in active chromatin in at least one tissue (FDR <5%; Fig. 1a; Supplementary Table S2). These data suggest variability across tissues: aorta, brain anterior caudate, and adipose are the most “permissive” tissues, while right atrium and spleen do not show any significant TE enrichment in active regions (Fig. 1a).

**Figure 1.**
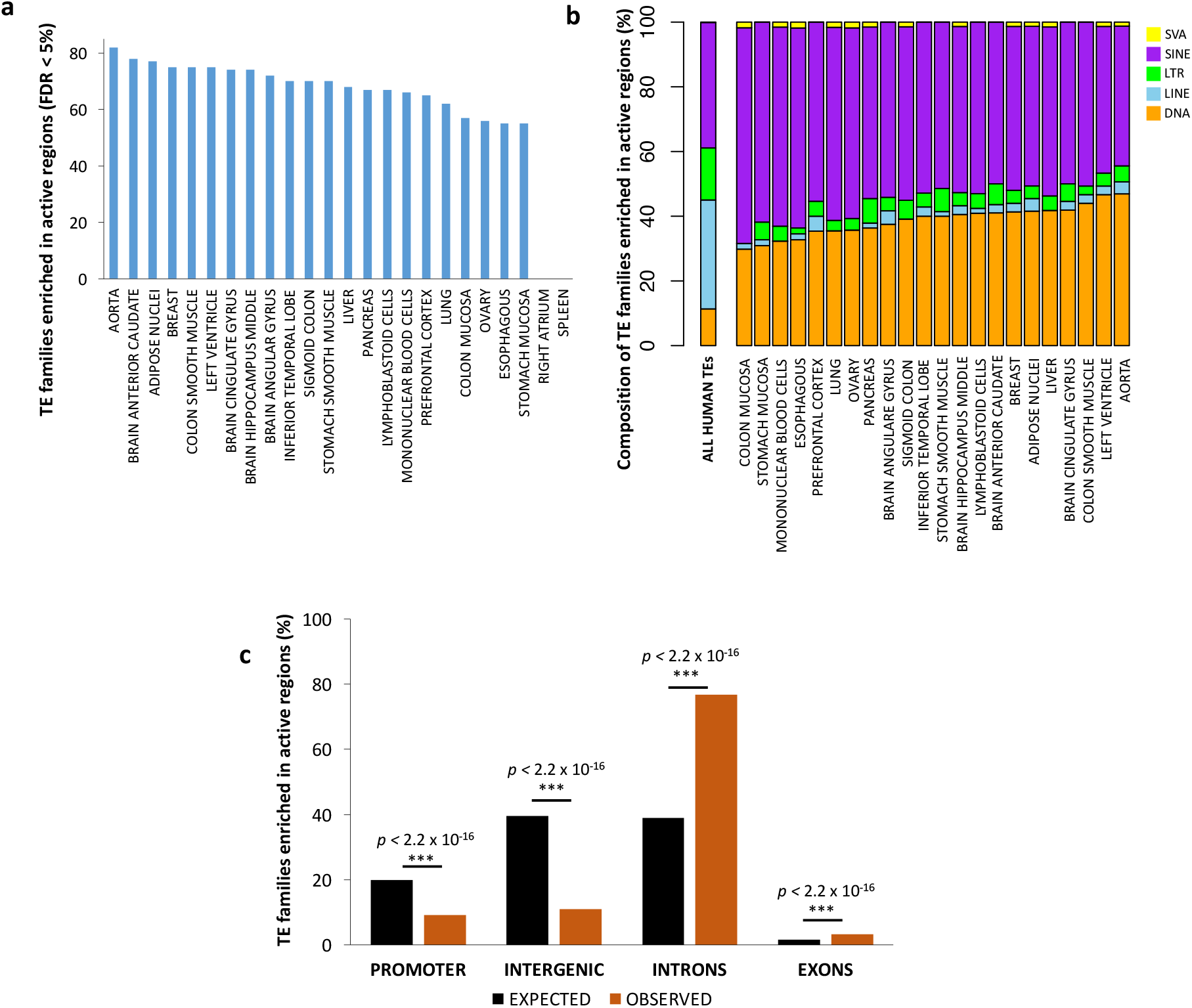
Transposable elements are enriched in active genomic regions. (A) The plot displays the numbers of enriched TE families in the active genomic regions for each tissue (FDR <5%). The distribution suggests a tissue-specific pattern. (B) Stacked-bar charts show TE class composition for the TE families enriched in active regions (FDR <5%). SINE and DNA transposons are the dominant TEs enriched in active regions. (C) The TEs enriched in active regions are depleted from promoters and intergenic regions, while they are significantly enriched in intronic regions.

SINEs and “cut and paste” DNA transposons are the classes most frequently enriched in active chromatin (Fig. 1b). SINE families, the most abundant human TEs (38.8% of the total), correspond to 43-66% of the TEs enriched in active regions (FDR < 5%), these fractions being more than expected by chance in all tissues (Proportion Test *p* < 2.2 × 10^−16^ for each tested tissue). Similarly, DNA TEs, that account for 11.3% of the annotated TEs, represent 29–47% of the transposons enriched in active regions (Proportion Test *p* < 2.2 × 10^−16^ for each tested tissue). In general, SINE-Alu elements are the most commonly enriched TEs (Supplementary Table S2).

Conversely, LTRs and LINEs are significantly depleted from active genomic regions of all tissues (Proportion Test *p* < 2.2 × 10^−16^ for each tested tissue; Fig. 1b). Finally, SINE-VNTR-Alus (SVAs), which are the least abundant TEs in the human genome (0.12% of the total annotate TEs in the human genome), are significantly overrepresented in active chromatin in 13/24 tissues; Fig. 1b).

We set out to investigate the TEs overlapping active regions. These TEs are depleted in active promoters and intergenic regions, but significantly enriched within active regions inside gene bodies, and in particular in introns (Fisher’s Exact Test *p-values* in Fig. 1c). More specifically, 96.3% of TEs enriched in gene bodies overlap introns, in line with the normally observed distribution of introns and exons in the human genome (Fig. 1c, Fisher’s Exact Test *p* > 0.05). We speculate that genomic regions containing active genes are more frequently accessible, thus providing a substrate for TEs to insert. Moreover, TEs present in the bodies of active genes may be less likely to be silenced than TEs in intergenic regions.

Using the same approach previously described for the active regions, we searched for TEs enriched in repressed genomic regions. Overall, 314 human TE families (25.4%) are significantly enriched in repressed regions of the genome in at least one tissue (FDR <5%; Fig. 2a; Supplementary Table S3). LTRs (predominantly ERV1) represent the large majority of the repressed TEs (Fig. 2b), followed by LINEs (predominantly L1s) and DNA TEs. Notably, ERV LTRs and L1 LINEs are among the most active TEs in the genome, and also have their own regulatory architecture [40, 41]. We thus surmise that these autonomous active TEs may be preferential targets of repressive marks.

**Figure 2.**
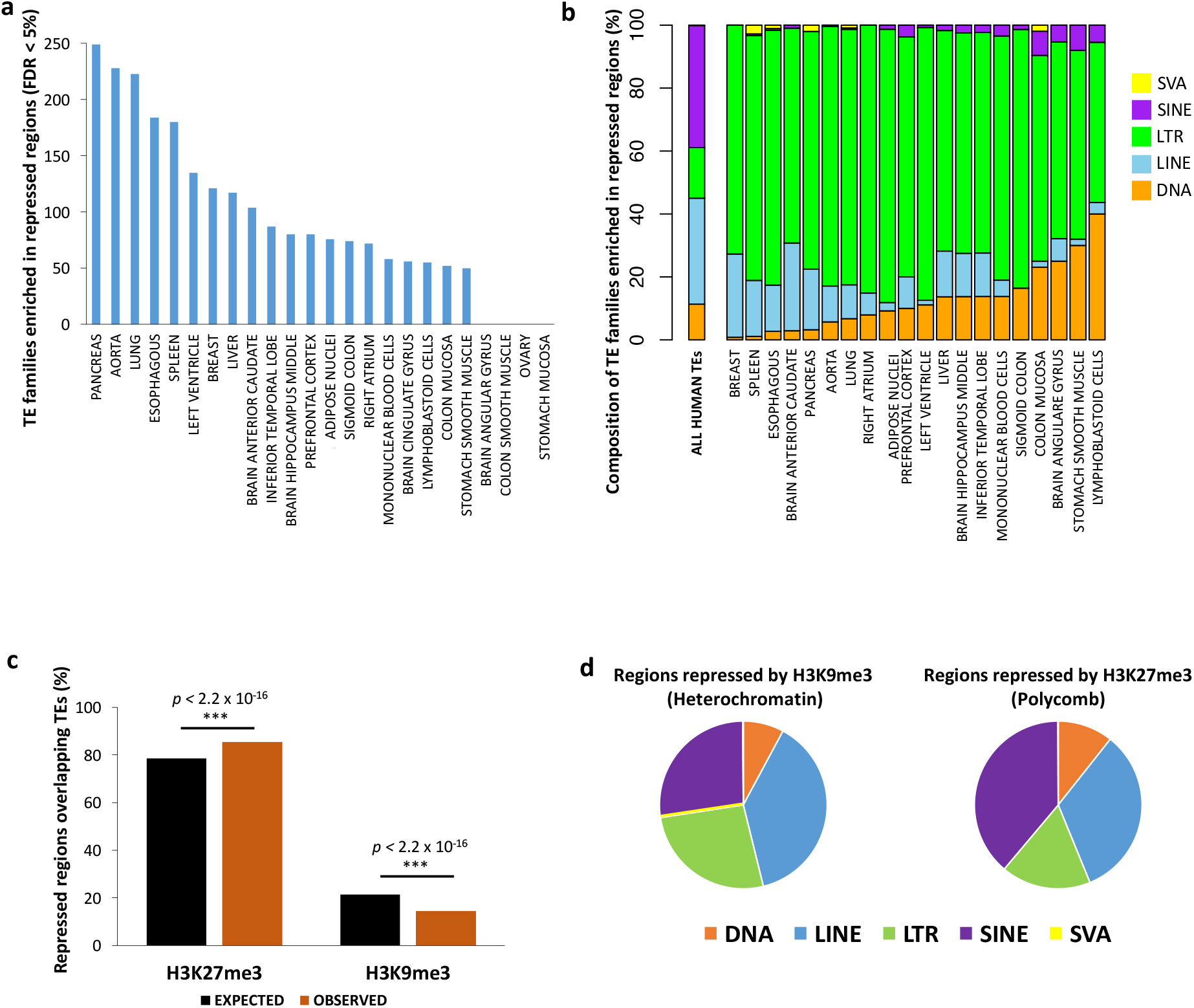
Transposable elements are enriched in repressed genomic regions. (A) The plot displays the numbers of enriched TE families in the repressed genomic regions for each tissue (FDR <5%). The distribution suggests a tissue-specific pattern. (B) Stacked-chart plot shows class composition for the TE families enriched in repressed regions (FDR <5%). (C) Across tissues, the repressed TEs overlap H3K27me3 more than expected by chance, while H3K9me3 is underrepresented. (D) Pie-charts show class composition for the TEs overlapping H3K27me3 and H3K9me3.

We note a very high variability in the number TE families enriched in repressed regions across tissues (Fig. 2a), as well as large differences in the composition of enriched TE classes in the repressed regions. Notably, the tissues that harbor the highest number of TE families enriched in repressed regions (pancreas, aorta, lung, spleen, esophagus, breast, and liver; Fig. 2a) are also those displaying the highest numbers of enriched LINEs in the same repressed regions (Fig. 2b).

### Different TE repression patterns in the human genome

We examined whether TEs preferentially overlap regions repressed via Polycomb Repressive Complex (H3K27me3) or via Heterochromatin (H3K9me3). Overall, 78.6% of the regions classified as repressed in the human genome across all tissues are bound by H3K27me3 (Polycomb Repressive Complex), while 21.4% are marked by H3K9me3 (Heterochromatin conformation). However, when we restrict the analysis to the repressed regions containing a TE, we report an overall higher than expected overlap with H3K27me3 (median across tissues 85.5%; Proportion Test across tissues *p* < 2.2 × 10^−16^; Supplementary Table S4; Fig. 2C), and a consequent underrepresentation of H3K9me3 (median 15.5%; Supplementary Table S4; Proportion Test *p* < 2.2 × 10^−16^; Fig. 2d). In 20/24 of the tested tissues, TEs are marked by H3K27me3 more than expected by chance (Proportion Test *p* < 2.2 × 10^−16^ for each of the 20 significant tissues; Supplementary Table S4). In the remaining four tissues this histone mark is instead underrepresented, while H3K9me3 is overrepresented: breast (H3K27me3 = 76.4%; Supplementary Table S4; Proportion Test *p* < 2.2 × 10^−16^), aorta (55.1%; Supplementary Table S3; *p* < 2.2 × 10^−16^), lung (48.9%; Supplementary Table S4; *p* < 2.2 × 10^−16^), and spleen (26.5%; Supplementary Table S3; *p* < 2.2 × 10^−16^). Notably, in these four tissues we detect the highest numbers of TE families enriched in repressed regions (Fig. 2a), and the highest proportion of repressed LINEs. We speculate that the heterochromatin state (H3K9me3) may be employed to target specific TE classes and families in a context specific manner [36].

We therefore tested whether different TE classes correlate with either heterochromatin (H3K9me3) or with Polycomb repressed chromatin (H3K27me3). LTRs, LINEs, and SVAs are overrepresented in regions marked by H3K9me3 (Fisher’s Exact Test *p* < 2.2 × 10^−16^; Fig. 2d). Conversely, SINEs and DNA TEs are significantly more likely to overlap H3K27me3 than expected by chance (Fisher’s Exact Test *p* < 2.2 × 10^−16^; Fig. 2d). Notably, SVAs are depleted from the regions marked by H3K27me3 (Fig. 2d).

These findings are consistent with recent reports suggesting that H3K27me3 and H3K9me3 target different transposon types in embryonic stem cells [42], and with a study reporting that LINEs, LTRs, and SVAs are the most abundant TEs repressed by H3K9me3 in induced pluripotent stem cells [42].

### Ancient TEs are enriched in active regions, while young TEs are repressed

We clustered the annotated human TEs in 35 age classes as in ref. 39 (e.g. Eutheria, Primates, Hominidae; Supplemental Table S6), and used the TE-Analysis shuffling script to test for enrichment of each age class in a given set of regions (see Methods). Using this approach, we assessed the age of TEs enriched in active and repressed genomic regions. Ancient TE classes (i.e. age classes older than the Eutheria lineage) are enriched in the active regions of all tested tissues (FDR <5%; Supplemental Table S6). These TEs are largely vertebrate or mammalian specific (Supplemental Table S6). Notably, the only tissues with an enrichment of young TEs (specifically primate specific) are blood related (Mononuclear and Lymphoblastoid Cells). These results are in agreement with an elegant study that discovered a key role of primate specific TEs in the regulatory evolution of immune response [25]. TE families enriched in active regions across at least 20 of the 24 tissues correspond to DNA TEs and SINEs (Supplemental Table S2). Despite a lack of enrichment of all young TEs taken together in active regions, 24 *Alu* families are in fact enriched in active regions.

In contrast, young TEs (i.e. TE classes younger than the Eutheria lineage split) are significantly enriched in the repressed regions of most tissues. In particular human specific TEs are enriched in the repressed regions of all brain related tissues (FDR <5%; Supplemental Table S6). These young TEs correspond to ERV LTRs, L1 LINEs, and SVAs, but only one family is found enriched in at least 20 tissues (MER52A), which is in line with the broad cross-tissue variability of the TEs enriched in repressed chromatin regions (see above).

Collectively, these data suggest that young TEs are predominantly silenced, while the older TE fragments still detectable in the human genome are now more tolerated.

### TE insertions are associated with gene expression variance across tissues

We employed GTEx data to test if TE insertions affect local gene expression. For this purpose, we first assigned each TE overlapping an active genomic region to its nearest gene transcription start site (TSS). Next, we divided all human genes in four categories (Supplemental Table S7): 1) Genes associated with TEs that are only found in active regions across tissues; 2) Genes associated with TEs that are found in active or repressed regions in a tissue-specific fashion; 3) Genes associated with TEs that are only found in repressed regions; 4) Genes never associated with TE insertions. Based on this classification, genes associated with a TE insertion in regions that are active in at least one tissue are characterized by significantly higher expression variance (normalized by mean expression) than genes either associated to repressed TEs or not associated to a TE (Wilcoxon’s Rank Sum Test *p* < 2.2 × 10^16^; Fig. 3). Similarly, the genes associated with TEs exclusively found in active regions have significantly higher expression variance than the genes associated with TEs present in both active and repressed regions (Wilcoxon’s Rank Sum Test *p* = 9.91 × 10^−8^; Fig. 3). We reasoned that TE insertions may happen more likely at longer genes located in gene deserts. However, even after correcting our model for gene density and gene length, the gene expression variance is still positively correlated with TE insertion in active regions (linear regression *p* < 2.2 × 10^−16^).

**Figure 3.**
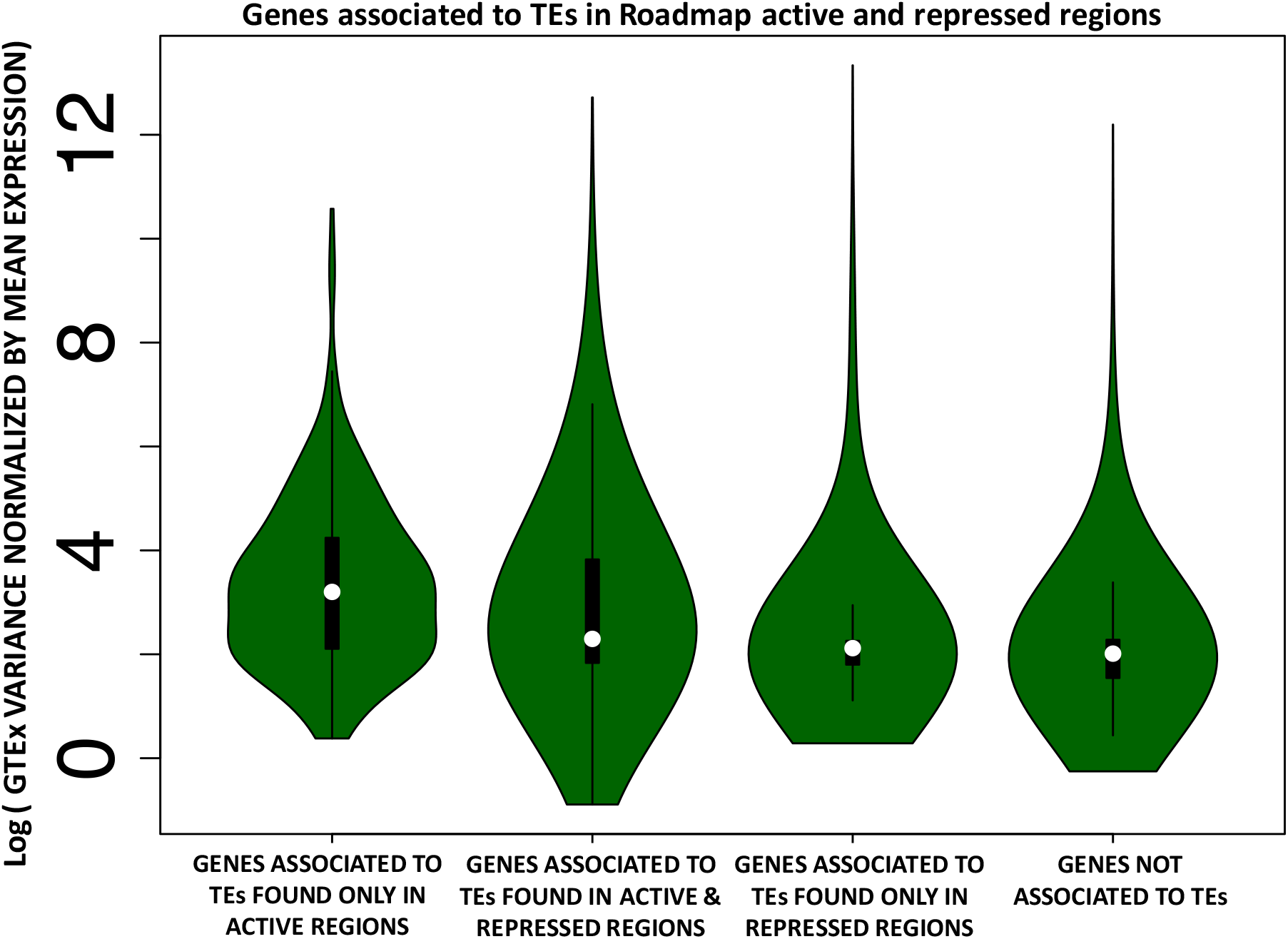
Genes with higher expression variance are more tolerant towards TE expression. Human genes were split into four categories: 1) Genes associated with TEs that are only found in active regions across tissues; 2) Genes associated with TEs that are found in active or repressed regions in a tissue-specific fashion; 3) Genes associated with TEs that are only found in repressed regions; 4) Genes never associated with TE insertions. The violin plots display the distribution of the GTEx gene expression variance, normalized by mean expression, for each of the four categories.

Together, these findings suggest that genes with local TEs overlapping active chromatin have higher variability in gene expression across tissues, and that genes consistently expressed across tissues (e.g. housekeeping and other essential genes) may be less tolerant towards TE insertions in their regulatory regions.

### Tissue-specific TE enrichment in active regions correlates with tissue-specific gene expression

We compared the relative enrichment in active regions of each TE family across tissues. Specifically, for each TE enriched in active regions (FDR < 5%), we leveraged the Odd Ratios from the permutation test of the TE-Analysis pipeline to compute Z-scores (i.e. effect sizes; see methods), and compare them across tissues. We find that TE enrichment varies substantially across tissues (Supplemental Table S5; Fig. 4), and many TEs exhibit tissue-specific enrichment in active chromatin (Fig. 4). For example, HERV15 (LTR) is significantly more enriched in the liver and in the stomach mucosa compared to any other tissue (Fig. 4). Motif analysis revealed that the liver regions of active histone modification overlapping HERV15 are enriched in motifs for EOMES (Supplemental File S2). This transcription factor (TF) has a key role in the hepatic immune response, instructing the development of two distinct natural killer cell lineages specific to this tissue [43]. Moreover, EOMES is also an established tumor suppressor in Hepatocellular Carcinoma [44]. Notably, HERV15 was recovered as significantly enriched in the human liver enhancers also in our previous study [33], suggesting that the findings of the present analysis are not likely to represent batch-specific effects of the Roadmap data.

**Figure 4.**
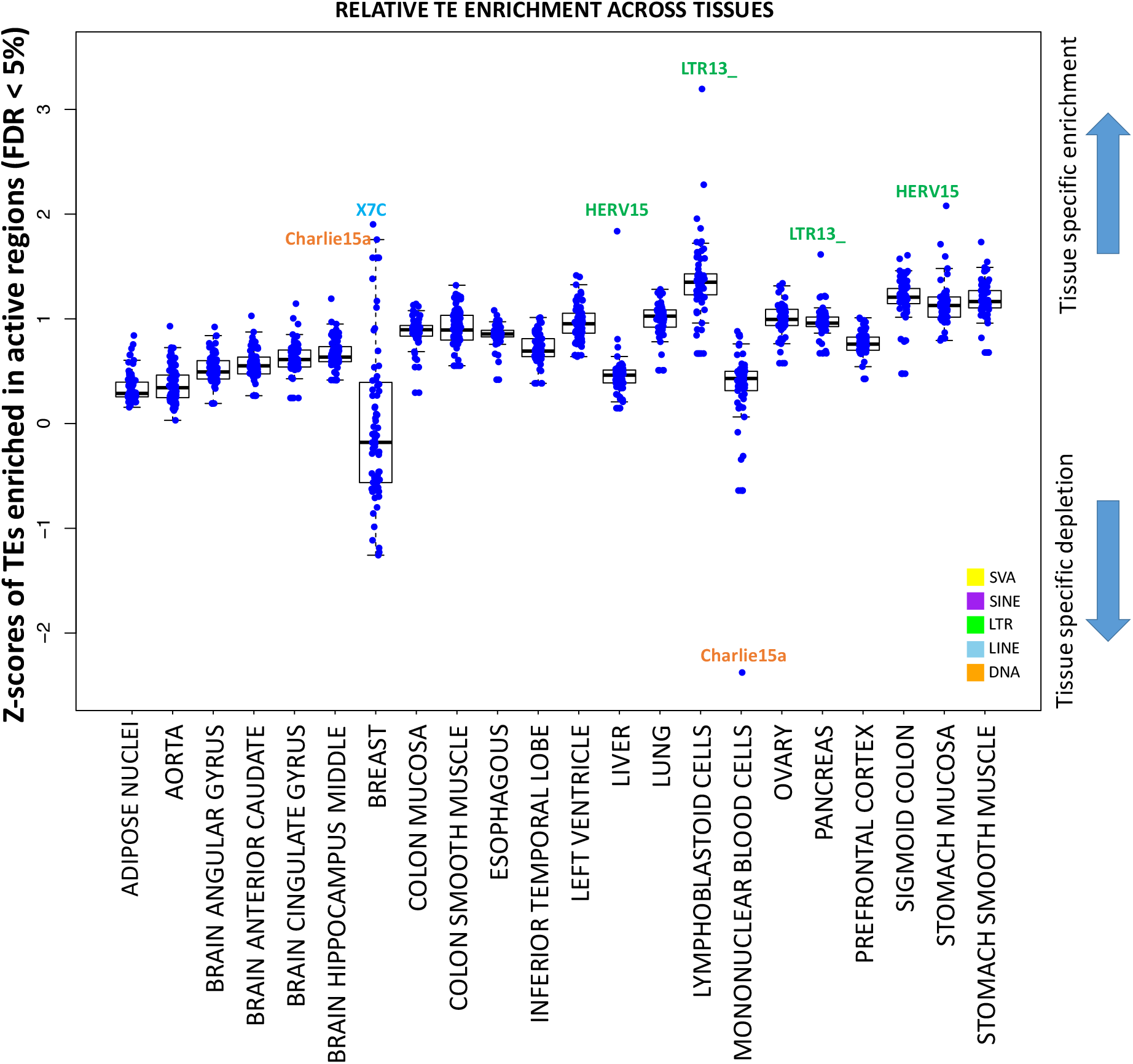
Transposable elements have tissue-specific enrichment in active regions. The plot displays the distribution of the effect sizes (Z-scores from permutation test, see methods) for each TE enriched in active regions (FDR < 5%), in each tissue. The higher the Z-score, the more tissue-specific is the enrichment.

Similarly, X7C (LINE) and Charlie15a (DNA TE), are the most enriched TEs within regions bearing active chromatin state in the breast. In the sequence of these we find enrichment for binding sites for key breast TFs as KLF5 and CPEB1 (Fig. 5a; Supplemental File S2). Notably, KLF5 is an essential regulator of hormonal signaling and breast cancer development [45], and is considered a breast cancer suppressor [46]. Similarly, CPEB1 mediates epithelial-to-mesenchyme transition in breast, and mice depleted of this gene showed increased breast cancer metastatic potential [47]. Interestingly Charlie15a shows tissues-specific depletion in the mononuclear blood cells (Fig. 4), highlighting a potential tissue-specific regulatory activity.

**Figure 5.**
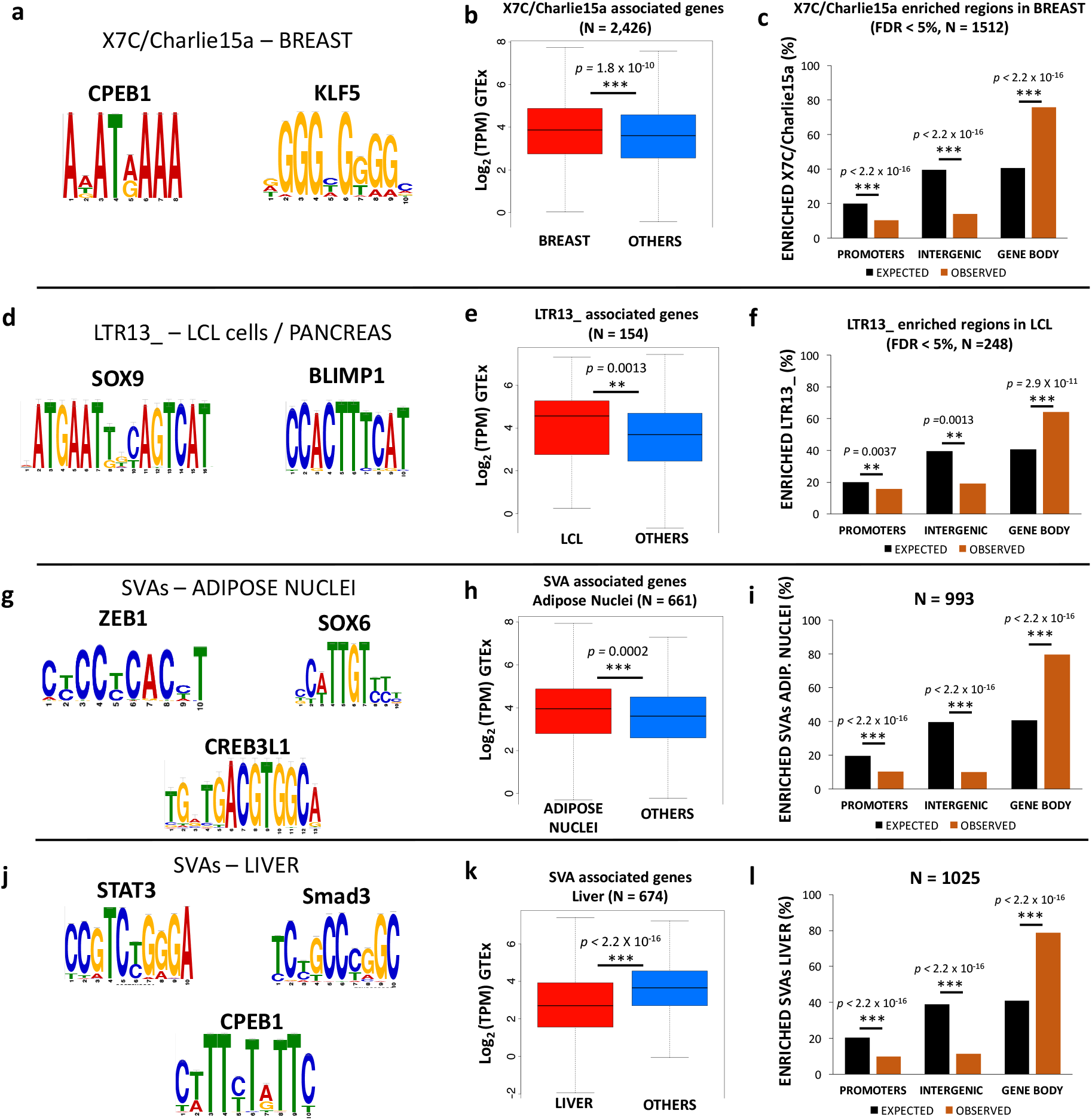
Tissue-specific TEs are enriched for TF binding sites, are mostly intronic, and affect gene expression. (a) Motifs enriched in the regions overlapping X7C and and Charlie15a TEs in the breast, (b) Boxplot comparing mean expression for the genes associated to X7C and and Charlie15a in the breast vs all the other tissues, (c) Genomic distribution of the regions overlapping X7C and and Charlie 15a TEs in the breast, (d) Motifs enriched in the regions overlapping LTR13_ TEs in pancreas and LCL cells, (e) Boxplot comparing mean expression for the genes associated to LTR13_ in the LCLs vs all the other tissues, (f) Genomic distribution of the regions overlapping LTR13_ in the LCLs. (g) Motifs enriched in the regions overlapping SVAs in the adipose nuclei, (h) Boxplot comparing mean expression for the genes associated to SVAs in the adipose nuclei vs all the other tissues, (i) Genomic distribution of the regions overlapping SVAs in the adipose nuclei, (j) Motifs enriched in the regions overlapping SVAs in the liver, (k) Boxplot comparing mean expression for the genes associated to SVAs in the liver vs all the other tissues. (I) Genomic distribution of the regions overlapping SVAs in the liver.

To assess the robustness of the enrichment of X7C and Charlie15a in the breast, we ran the TE-Analysis pipeline on publicly available H3K27ac and H3K4me1 data generated by Encode from the breast epithelium and from the MCF7 cell line [48]. Notably, these two TEs were also significantly enriched in the Encode data (FDR < 5%), suggesting that batch effects are unlikely strong drivers of this trend.

Analogously, LTR13_ is the most enriched TE in the active chromatin of pancreas and Lymphoblastoid Cell Line (LCL). These LTR copies are enriched for binding sites for SOX9 and PRDM1/Blimp-1 (Fig. 5d; Supplemental File S2). SOX9 is a master regulator of the pancreatic program [49], while PRDM1/Blimp-1 has a central role in determining and shaping the secretory arm of mature B Lymphocyte differentiation [50].

We next tested whether tissue-specific TE enrichment in active chromatin (Fig. 4, 5a–f) correlates with tissue-specific-changes in gene expression. Specifically, we tested the TE families showing the highest degree of tissue-specific enrichment (Fig. 4: HERV15/liver, LTR13_/LCL, X7C-Charlie15a/breast). With the exception of HERV15/liver (Wilcoxon’s Rank Sum Test *p* > 0.05), in the other tested instances (LTR13_/LCL; X7C-Charlie15a/breast) the tissue-specific enrichment of the TEs in active chromatin regions is associated with a significant change in the associated gene expression (Wilcoxon’s Rank Sum Test *p-values* in Figs. 5b,e). These findings support a possible regulatory role for the co-opted TEs.

To better understand how these tissue-specific TEs may be involved in the regulation of gene expression, we investigated what typology of genomic region they overlap (i.e. promoter, intergenic, introns, exons). Both X7C/Charlie15a in breast and LTR13_ in LCLs are significantly depleted in promoter and intergenic regions, but overrepresented in gene bodies (Figs. 5c, f), 97.8% (X7C/Charlie15a) and 96.4% (LTR13_) of them respectively found in introns.

The Roadmap data did not include H3K27ac profiles for all tissues. Therefore, to further characterize these intronic regions, we leveraged again the publicly available H3K27ac and H3K4me1 Encode data for the breast (Breast epithelium and MCF7 cell line; [48]). These data reveal that 57.0% of the intronic regions containing X7C or Charlie15a overlap a H3K27ac or H3K4me1 peak, thus suggesting that most of these regions likely represent breast intronic enhancers. As comparison, only 33.7% of random intronic regions of the same size and number of the ones overlapping X7C/Charlie15a TEs are overlap a H3K27ac or H3K4me1 peak (Fisher’s Exact Test *p* < 2.2 × 10^−16^).

Collectively, these findings point towards a model in which specific TE families, largely belonging to LTR (ERVs) and DNA TE classes, have more regulatory potential than other transposons. Furthermore, our data expand upon previous findings suggesting that ERVs that escape repression can have a significant impact on the host gene regulation [9, 25, 26, 33, 51, 52].

### SVAs exhibit tissue-specific regulatory activity

In our recent work, we demonstrated that a large fraction of human specific cis-regulatory elements in the liver are SVA transposons, which typically function as transcriptional repressors, at least in this tissue [33]. SVAs are very young transposons, being Hominidae (SVA_A, B, C and D) and human specific (SVA_E and F). According to Roadmap data, SVAs are enriched in the active regions of 13/25 tissues (Fig. 1b), and mainly corresponded to SVA_A copies (Supplementary Table S4). We first assessed the potential contribution of SVAs to gene regulation of two of these tissues: the adipose nuclei and the liver.

In both tissues, SVAs provide binding sites for key transcription factors (Fig. 5g, j; Supplemental File S2). ZEB1 is the master regulator of adipogenesis [53, 54], and, based on GTEx data, is ten times more highly expressed in adipose tissue compared to the liver. Similarly, SOX6 contributes to the developmental origin of obesity by promoting adipogenesis, and has a key role in adipocyte differentiation [55]. Consistent with the data reported for other tissues, SVAs associated with active chromatin in adipose nuclei and liver are strongly enriched in gene bodies (Figs. 5i, l). Genes associated with SVAs in the adipose nuclei are significantly more highly expressed in this tissue compared to other tissues (Wilcoxon’s Rank Sum Test *p* = 0.0002; Fig. 5h), suggesting that SVA elements can work as transcriptional activators, at least in the adipose tissue.

In the liver, SVAs in active regions are enriched for hepatic regulators like CPEB1, that mediates insulin signaling in the liver (Fig. 5j; [56]), and STAT3, that regulates liver regeneration and immune response and negatively modulates insulin action (Fig. 5j; [57]). However, the liver SVAs are also enriched for established transcriptional repressors, like Smad3 (Fig. 5j). Consistently, genes associated with liver active SVAs exhibit lower expression in this tissue compared to all the others (Wilcoxon’s Rank Sum Test *p* < 2.2 × 10^−16^; Fig. 5k), supporting the previously proposed repressive role of SVAs in the hepatic system [33].

### Discussion

The contribution of transposable elements (TEs) to gene regulation was proposed over half a century ago [10–13] and considerably expanded over the last two decades, largely due to the advances in next generation sequencing [14–36].

In order to gain insights in this topic, we identified TEs enriched in active and repressed genomic regions of 24 human tissues, using Roadmap and GTEx data. Our analyses provide a novel integrated overview of the potential impact of TEs to the human gene regulation across multiple tissues, correlating the enrichment of TE copies in active chromatin to tissue-specific gene expression. In fact, many of the previous studies have proposed that TEs are frequently enriched in cis-regulatory elements and lncRNAs [21, 22, 33, 39, 58], but the actual effect of the presence of TEs on the associated gene expression was not tested on a large scale.

Recent work has evaluated the prevalence of TE-derived DNA in enhancers and promoters across mouse cell lines and primary tissues [35]. The present study builds upon this by investigating the dynamics of TE recruitment and the potential effects on tissue-specific gene expression.

We demonstrate that ~10% of the TEs identified in the human genome are significantly enriched in active regions (promoters, intergenic enhancers, intronic enhancers) of 24 different human tissues. In general, we report a high degree of variability of TE enrichment in the active and repressed genome across tissues, and detect multiple instances of TEs displaying potential tissue-specific regulatory function. We acknowledge that the correlation between tissue-specific TE enrichment in active regions and the tissue-specific changes in gene expression does not necessarily underly a causal role for the TEs. On the other hand, while it is possible that the changes in gene expression are simply due to the presence of a tissue-specific active histone mark, we also find that in all of the tested cases the enriched TE sequence provides binding sites for transcription factors that are master regulators for that specific tissue. This is consistent with the changes in the gene expression of associated (i.e. adjacent) genes and could explain why these TE insertions are retained by selection.

Enriched TEs are typically distributed along gene bodies, likely functioning as intronic enhancers. We reason that this may be explained by the assumption that TEs located within intra-genic regions are less likely to be repressed or removed. In agreement with these findings, a recent study has shown that TEs are depleted in human promoters and intergenic enhancers across multiple tissues [35]. In this context, we see a correlation between gene expression variance and the insertion of TEs in their loci or regulatory regions. This may suggest that genes consistently expressed across tissues are less prone towards TE co-option in their regulatory networks, but future analyses in this direction will be needed to further characterize this phenomenon.

On the other hand, L1 LINEs and ERV LTRs are the most frequently enriched TE classes in the repressed regions. L1 retrotransposons are among the most active TEs in the human genome [59], and several studies have demonstrated that they are also active in brain tissues (e.g. hippocampus), and can contribute to neuronal genetic diversity in mammals [60–63]. Both L1s and LTRs possess their own regulatory architecture, and we speculate that their preferential silencing prevents these TEs from interfering with gene regulatory networks. Despite this, we demonstrate that LTRs that escape repression may be co-opted in a tissue-specific manner in the active regulatory regions, putatively as a consequence of their regulatory potential.

We show that TEs enriched in repressed regions of most tissues are generally young, while TEs enriched in active regions of most tissues generally predate the split of eutherian mammals. This is consistent with an accumulation of mutations in these ancient copies that would have increased the likelihood to generate binding sites for transcription factors, and thus the probability for the TE to be co-opted in the regulatory networks. An alternative explanation could be that young TE insertions in active chromatin regions are more likely to be removed by purifying selection than the new insertions in repressed regions, since the latter are more likely to have a neutral impact.

Finally, we demonstrate that SVAs, previously characterized as transcriptional repressors in select cell-types [33, 64], can act as both activators or repressors in a tissue-specific fashion.

## Conclusions

In summary, we present a comprehensive overview of the contribution of TE copies to human gene regulation: not only do they provide an important source of evolutionary novelty for the genome, but they can also function with tissue-specific patterns, modulating the expression of key genes and pathways.

## Methods

### TE-Analysis pipeline

To test for TE enrichment in active and repressed regions, we used the TE-Analysis pipeline v 4.6 ([39]; https://github.com/4ureliek/TEanalysis). This pipeline is designed to output the TE composition of given features, such as TE counts and TE amounts, aiming to detect potential TE enrichments in the select features. Roadmap annotated BED files (i.e. files listing the coordinates of annotated genomic regions) for each of the 24 tissues were downloaded (http://egg2.wustl.edu/roadmap/data/byFileType/chromhmmSegmentations/ChmmModels/coreMarks/jointModel/final/; last access: 10/4/2017). One file per tissue was downloaded (TISSUE_ID_coreMarks_dense.bed.gz”; Supplementary Table S1). From each of the 24 BED files, we produced two different files: one for the regions enriched with epigenomics hallmarks of active chromatin (hereafter “active regions”. Histone marks: H3K4me1, H3K36me3, H3K4me3. Roadmap annotations: “TssA”, “TssAFlnk”, “TxFlnk”, “Tx”, “TxWk”, “EnhG”, “Enh”, “TssBiv”, “EnhBiv”), and one for the regions with signature of repressed chromatin (hereafter: “repressed regions”. Histone marks: H3K27me3, H3K9me3. Roadmap annotations: “Het”, “ReprPC”, “ReprPCWk”).

For each tissue, we tested for TE enrichment in the “active” and “repressed” BED files using the “TE-analysis_Shuffle_bed.pl” script v 4.3. Specifically, this script assesses which TEs are significantly enriched in a set of features (BED files) by comparing observed overlaps with the average of *N* expected overlaps (here 1000). These expected overlaps were obtained by shuffling the genomic position of TEs. TE annotations were downloaded from the University of California Santa Cruz Genome Browser (RepeatMasker, Hg19 version; [65]).

The “TE-analysis_Shuffle_bed.pl” script was run with Bedtools v2.27.1 [66] and the following parameters:

-f Roadmap_BEDFILE (active or repressed)
-q RepeatMasker.out (TE file, hg19)
-n 1000 (number of bootstrap replicates)
-r hg19.chrom.sizes
-g 20141105_hg38_TEage_with-nonTE.txt (distributed with the pipeline)
-s rm (shuffles the TEs within their genomics position)

The script performs a two-tailed permutation test to assess the enrichment (or depletion) of each annotated TE in the given regions (Roadmap regions), thus assigning a *p-value* to each annotated TE. Additionally, we corrected for multiple testing by applying a False Discovery Rate (FDR; [67]). Only TEs with FDR < 5% were retained, considered significantly enriched in the given tissue, and used for downstream analyses.

### Composition of enriched TEs

To characterize TEs enriched within active and repressed regions of each tissue (e.g. Figs. 1b, 2b), each TE was assigned to one of the major TE classes: DNA transposons, LINEs, LTRs, SINEs, SVAs, according to RepeatMasker annotations. To assess the genomic distribution of the enriched TEs (e.g. Figs. 1c, 2c), we considered as 1) PROMOTERS: all of the regions found within +/- 1 Kb from an annotated TSS (Gencode_v19 comprehensive annotations). 2) GENE BODIES: all of the regions overlapping an annotated gene but not overlapping the promoter region. 3) INTERGENIC - all of the regions not overlapping an annotated gene and distant > 1 Kb from a TSS.

### Correlation between TE insertion and variance in gene expression

We calculated the variance and mean of the TPM (Transcripts Per Million) for each gene using GTEx data. We assigned each TE overlapping an active or a repressed region to the closest gene, based on the distance to the nearest transcription start site. Next, we divided all human genes in four categories: 1) Genes associated with TEs that are only found in active regions across tissues; 2) Genes associated with TEs that are found in active or repressed regions in a tissue-specific fashion; 3) Genes associated with TEs that are only found in repressed regions; 4) Genes never associated with TE insertions. Gene expression variance, normalized by mean expression, was compared across the four categories. Gene density and gene length were used as covariates for the model. Specifically, gene density was calculated as the amount of exonic sequence present within +/- 100 Kb from each gene. In summary, the following model was used:

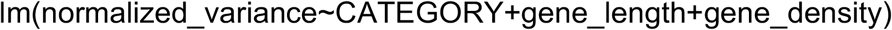

Variance was normalized by average expression across tissues.

### Computation of Z-scores for tissue-specificity

For each TE enriched in active regions (FDR < 5%), we used the Odd Ratios (OR) from the permutation test of the TE-Analysis pipeline to compute Z-scores with the following equation: (OR – mean(OR)) / sd(OR). Z-scores can be found in Supplemental Table S5.

### Motif analyses

Motif analyses were performed using the Meme-Suite [68], and specifically with the Meme-ChIP application. Fasta files of the regions of interest were produced using BEDTools v2.27.1. Shuffled input sequences were used as background. *E-values* < 0.001 were used as threshold for significance [68].

### Testing for TE co-option on gene expression

For each human gene and for each tissue, GTEx provides the mean of the TPMs (Transcripts Per Million). To test whether tissue-specific TE enrichment correlates with tissue-specific changes in gene expression, for each gene associated with a TE of interest, we used the mean TPMs to compare the expression of genes in the tissue of enrichment Vs the average of the gene expression of the same genes in all the other considered tissues (i.e. mean of TPMs across all the other tissues).

### Statistical and genomic analyses

All statistical analyses were performed using R v3.4.1 [69]. Figures were made with the package ggplot2 [70]. BEDTools v2.27.1 was used for all the genomic analyses.

## Acknowledgements

We thank Roadmap and GTEx Consortia for the generation of invaluable data. MT thanks his current P.I. (Alessandro Gardini, The Wistar Institute) who granted him time and freedom to work on this project. We also thank the two anonymous reviewers for their valuable suggestions and insights.

## Authors’ contributions

MT and CDB designed the project. MT, AK, and CDB analyzed the data. MT, AK, and CDB wrote and approved the manuscript.

## Ethics approval and consent to participate

N/A

## Consent for publication

N/A

## Availability of data and material

N/A

